# Learning a force field from small-molecule crystal lattice predictions enables consistent sub-Angstrom protein-ligand docking

**DOI:** 10.1101/2020.09.06.285239

**Authors:** Hahnbeom Park, Guangfeng Zhou, Minkyung Baek, David Baker, Frank DiMaio

## Abstract

Accurate and rapid calculation of protein-small molecule interaction energies is critical for computational drug discovery. Because of the large chemical space spanned by drug-like molecules, classical force fields contain thousands of parameters describing atom-pair distance and torsional preferences; each parameter is typically optimized independently on simple representative molecules. Here we describe a new approach in which small-molecule force field parameters are jointly optimized guided by the rich source of information contained within thousands of available small molecule crystal structures. We optimize parameters by requiring that the experimentally determined molecular lattice arrangements have lower energy than all alternative lattice arrangements. Thousands of independent crystal lattice-prediction simulations were run on each of 1,386 small molecule crystal structures, and energy function parameters of an implicit solvent energy model were optimized so native crystal lattice arrangements had lowest energy. The resulting energy model was implemented in Rosetta, together with a rapid genetic algorithm docking method employing grid based scoring and receptor flexibility. The success rate of bound structure recapitulation in cross-docking on 1,112 complexes was improved by more than 10% over previously published methods, with solutions within <1 Å in over half of the cases. Our results demonstrate that small molecule crystal structures are a rich source of information for systematically improving computational drug discovery.

## INTRODUCTION

Classical force field parameterization based on liquid thermodynamic data and quantum chemistry typically proceeds by fitting different subsets of parameters on individual representative molecules independently^1–4^. A challenge with this approach is the *transferability* of the resulting model to systems not included in the parameterization set^5,6^. For example, bond torsional parameters are often obtained by computing the energies of a set of conformations of test molecules with quantum chemistry, and then subtracting the electrostatic and van-der-Waals contributions. However the resulting fitted function is highly dependent on the molecules selected for training. Using such a model to evaluate the energetics under different flanking chemical groups often yields inaccurate results^7,8^. Roos *et al*^3^ showed that this issue could be resolved by expanding to hundreds of thousands of parameters fit to reproduce quantum chemistry calculations on many thousands of small molecules. We hypothesized that a balanced and transferable energy model involving far few parameters could be learned by utilizing the many thousands of crystal structures of small molecules, which span a large diversity of chemical space^9–11^. Since these crystal structures form spontaneously, the majority of these must be very low free energy states, and hence the sum of the intra- and interatomic interaction energies must be low compared to almost all alternative packing arrangements and conformations of the molecule in the majority of cases.

The key ideas underlying our approach are: a) generation of large numbers of alternative “decoy” lattice packing and conformational arrangements of a set of small molecules with known crystal structures; and b) simultaneous optimization of a large set of force field parameters, such that the experimentally observed crystal structures have lower energies than all of the alternative states. The advantages of this approach are that parameters are obtained directly from structural data of molecules^10,12^ that are generally larger and more similar to drug-like compounds than the simple molecules traditionally used for QM calculations. Moreover, as the energy of a crystal involves tradeoffs between different forces, this approach should yield a balanced force field which can (for instance) accurately model the subtle interplay between deviations from bonded geometry minima and optimization of non-bonded interactions.

## METHODS AND MATERIALS

### Overview of the approach

We sought to develop a generalized force field for drug discovery following these three phases sequentially: i) to generate small-molecule docoy lattices, ii) to design and train a force field to discriminate native lattices from among these decoys, and iii) to validate our forcefield on small-molecule docking experiments. We first generated alternative packing arrangements for small molecules using a diverse set of 1,386 small molecule crystal structures from the Cambridge Structural Database (CSD)^10,12^ (870 for training and 516 for testing), by adapting Rosetta symmetry docking machinery^13^ to sample space groups, lattice parameters, rigid-body and internal conformation of each small molecule (**Fig 1a**). We simultaneously fit 175 non-bonded parameters for a generalized implicit solvent force field with 57 atom types (**Table S1**) plus 269 parameters for a torsion model conditioned on both constituent atom types and bond types^8^. The 444 free parameters were optimized to simultaneously maximize the energy gap between the experimentally observed lattice and the sampled alternative arrangements, and the fit to small molecule thermodynamic and protein-ligand complex structural data (**Fig 1b**) using the Simplex-search-based dualOptE algorithm^15^. Nine iterations of parameter optimization followed by crystal lattice regeneration were carried out; the final energy model is referred to as *RosettaGenFF. RosettaGenFF* was then tested on ligand docking benchmark sets using the newly developed docking tool Rosetta *GALigandDock*. In the following sections, we describe crystal lattice prediction protocol to generate training data, the energy model and parameter optimization procedure, and ligand docking method and dataset, in more detail.

**Fig 1.**
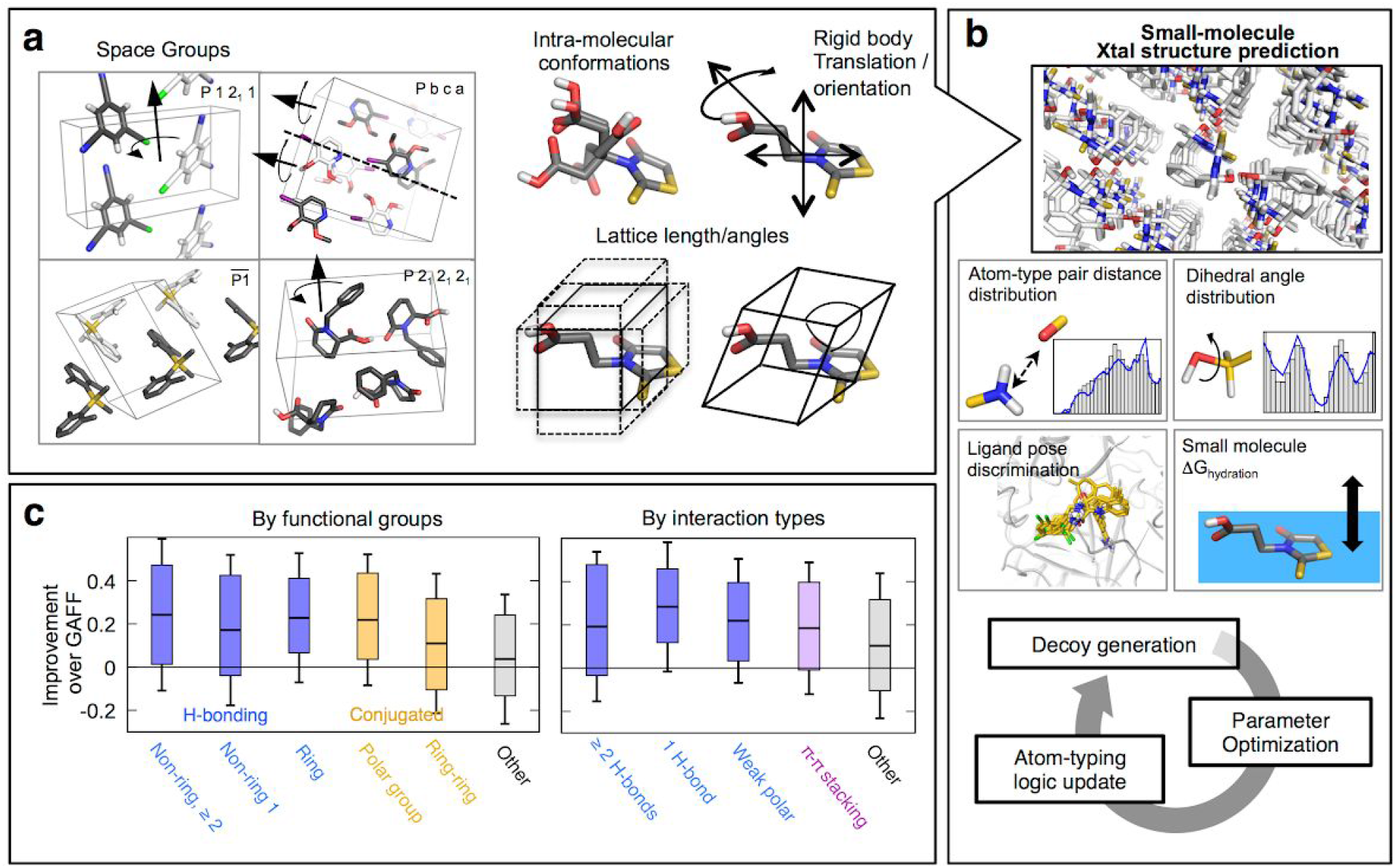
Force field optimization using small molecule crystal structures. **a)** Structure perturbation operations in Monte Carlo conformational search used for small molecule crystal structure prediction. Random space group assignment is done at the start of each simulation, followed by 50 cycles of interspersed lattice parameter and intramolecular perturbation followed by minimization over all degrees of freedom. **b)** Schematic overview of iterative parameter optimization procedure integrating small molecule crystal structure prediction, the KL divergence of sampled dihedral angle and distance distributions compared to reference distributions derived from ~4,000 small molecule crystal structures, ligand-protein docked pose discrimination tests on 215 complexes each containing hundreds of pre-sampled conformations^27^, and agreement with experimental hydration free energy for 643 small molecules^28^. At every iteration, new force field parameters are obtained by simplex optimization using dualOptE^15^, atom type classification logic is updated as necessary, and new low energy decoy lattice structures are generated. **c)** Comparison of performance against generalized Amber force field (GAFF^1^), decomposed by functional groups (left) or by interaction types across symmetry units (right). Statistics are collected from all molecules containing corresponding features and hence individual molecules can be counted multiple times.

### Crystal structure prediction protocol

We developed a lattice-docking protocol to sample small molecules in various crystallographic space groups. To handle space groups with mirror symmetries, Rosetta’s symmetry machinery^16^ was extended to allow mirror symmetry operations. For each space group, we expose as degrees of freedom (DOFs) the internal coordinates of the asymmetric unit, the rigid-body orientation of the molecule, and the dimensions of the lattice (**Fig 1a**). The symmetry machinery in Rosetta allows these DOFs to be sampled as well as minimized while maintaining the overall symmetry of the system.

Each run of structure prediction is carried out by running Metropolis Monte Carlo with minimization (MCM) search. At the beginning, lattice parameters are randomly assigned in a range of 0.2 to 1.0 on cell dimensions and 60 to 120 degrees on lattice angles. The input ligand conformation is also randomized by uniformly sampling all rotatable dihedral angles and rigid body placements in the lattice. Starting from this initial lattice, perturbation of one of the following sets of DOFs is attempted (**Fig 1a**) at each MCM cycle: i) translation or rotation of molecule, ii) a single dihedral angle in molecule, and iii) all lattice lengths or angles. Perturbation magnitudes are randomly selected from normal distributions with standard deviations of 0.5 Å / 2.5° / 5.0°, for translation / rotation / dihedral angles of ligands, respectively, and (0.5\**sgmultiplicity*) Å for lattice dimensions, where *sgmultiplicity* tries to capture the number of symmetric operators along each axis in a given spacegroup, and is generally larger for space groups with higher symmetry. Lattice angles are sampled by allowing the random axis moves to modify the crystal axis direction as well as its magnitude. Subsequent minimization is made simultaneously on *all* DOFs, and the Metropolis criteria is applied. The lowest energy conformation after 50 cycles is returned.

Training and validation sets of crystal structures were collected from the Cambridge Structural Database (CSD)^10,12^ satisfying the following conditions: (i) has one molecule per asymmetric unit; (ii) has solvent content less than 1%; (iii) is composed of only the elements H,C,N,O,S,P,F,Cl,Br,I; and (iv) has at least three and at most twelve rotatable bonds. We first curated an *extended training set* consisting of ~4,000 molecules and used for deriving torsion and distance statistics (**Fig 1b**). 870 molecules in the set were taken to generate decoys for training. A separate validation set of 516 molecules was later collected from the CSD (independent of the extended training set) with the same conditions mentioned above. For each small-molecule crystal lattice, thousands of structures are generated by repeating independent MCM structure predictions starting from random assignments of space group and ligand conformation. Initial ligand conformations were selected from among a pool of maximum 10 structures sampled by “confab” mode in *openbabel*^17^. The space group is randomly assigned amongst a list of most commonly observed ones in extended training set according to the chirality of the molecule: P 1 2_1_/c 1, P 1 2_1_/n 1, P-1, C 1 2/c 1, P b c a, P n a 2_1_, C 1 c 1, P b c n, P c a 2_1_, P c c n, and P 1 1 2 for achiral molecules; P 2_1_ 2_1_ 2_1_, P1 2_1_ 1, C 1 2 1, and P 2_1_ 2_1_ 2 for chiral molecules. In addition to these “decoy” structures, near-native conformations were added to the conformation pool by running the same protocol without initial randomization. A total of > 1,000 *de novo* predictions and >100 native perturbations were made for each molecule. An example command-line for performing crystal lattice prediction can be found in **Supplemental Data**.

### RosettaGenFF

The energy model presented in this study, hereafter *RosettaGenFF*, integrates two distinct “sub-models.” The first is the previously developed Rosetta protein energy model^15^, which is applied to any of the 20 canonical amino acids; more details can be found in Ref^15,21^.

Non-protein molecules and their interactions with canonical amino acids are described by a set of generic energy terms developed in this study:

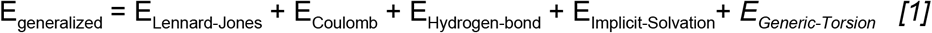

with atomic parameters defined for Lennard-Jones (LJ) and implicit solvation following the *generic atom types* (see below and also **Table S1**). As the partial charges used in Coulomb energy calculations are more molecular properties than atomic properties, we obtain them for each compound using AM1-BCC calculations^14^ and keep them fixed during model fitting. The functional forms of these terms are shared between the protein and generic sub-models. An exception is for describing torsion preference: in the protein energy model, for LJ and Coulombic interactions three or fewer bonds apart are ignored to avoid overlap with statistical torsion potentials, while in the generic energy model, only interactions one or two bonds apart are ignored.

#### Generic atom types

Our general strategy for assigning a distinct generalized atom type to each ligand atom is inspired from OPLS-AA force field^22^. We consider 35 common and unique functional groups containing at least one O,N,S,P in organic molecules listed in **Table S1**. When the atom does not belong to any of these functional groups, more general atom types are assigned by looking at element type and hybridization state (similar to Tripos force field^23^). Then the atom type is further specified based on the number of hydrogens attached in order to take into account variations in desolvation penalty, a unique aspect associated with implicit solvation energy model. The initial non-bonded parameters were determined by considering the “best matching” atom in Rosetta’s protein energy model^15,21^, followed by manual corrections on 9 LJ parameters to better reproduce experimental bulk liquid properties^24^. Note that atom types and their definitions were refined in between rounds of parameter optimization. The final list is given in **Table S1**.

#### Generalized torsion term

Our generalized torsion energy model follows the Karplus model, representing torsion potentials as a series of cosine functions up to 4^th^ order for an improved description of weakly conjugated systems^25^. Coefficients are assigned based on the atom types of the four constituent atoms and the bond order of the central bond. These parameters are optimized through the procedure following. First, the number of torsion occurrences are counted in the *extended training set* of small molecule crystal structures (see *Dataset* below). Torsion types observed at least 50 times were assigned unique torsion coefficients, yielding 150 torsion types. The remaining torsions are handled by atom-type grouping, with a total of 65 additional torsion classes. With 4^th^ order expansion of the Karplus equation, there are a total of 860 (=215 x 4) parameters. We further reduce this parameter set to 269 by restricting the coefficient order based on chemical intuition (e.g. torsions with strong preferences to planar conformations may only have non-zero first and second-order coefficients). The initial parameter set for optimization was brought from the best matching torsion in the OPLS-AA force field^22^.

### Parameter Optimization

Energy parameters were optimized by iteratively applying *dualOptE*^15^ primarily to maximize the energy gap between near-native and decoy lattices (**Fig 1b**). First, crystal lattice conformations were generated using the previously described lattice sampling method. Then *dualOptE* was run for 400 to 700 cycles of Nelder-Mead simplex minimization^26^, obtaining an optimal parameter set for the given atom type definition logic and decoy sets. The objective function used in dualOptE is represented as a weighted sum of metrics measuring performance on several specific tasks listed below. The number of parameters optimized at each dualOptE run ranged from 100 to 150, reduced from the 444 total parameters (269 torsional, 114 LJ, 57 solvation, and 4 hydrogen-bonding weight parameters) by grouping or sub-sampling parameters for efficiency. Finally, atom-type classification logic was updated by visually inspecting the failures originating from mistyping. This procedure -- from decoy generation to parameter optimization-was iterated 9 times until atom typing logic converged.

A first phase of optimization (the “*condensed phase*”) was carried out for the first 6 iterations. Here, LJ, hydrogen bonding, and torsion parameters are optimized considering two tasks: *lattice discrimination test* and *atomic geometry matching* (individual tasks are described below). During this phase, the solvation term was turned off, and electrostatics and hydrogen bonding terms were upweighted to their strength in a dielectric media with electrostatic permittivity of 2.0. A second phase (the “*solvent phase*”) was carried out for the final 3 iterations, beginning with parameters from the end of the condensed phase. Individual solvation and LJ parameters-together with a global weight controlling torsional energies -- were optimized simultaneously. Two additional tasks considering solvation energies were added to the overall optimization objective function (**Fig 1b**): *ligand pose discrimination* and *hydration free energy recapitulation*. These two tasks were critical for balancing components in the energy model. The ligand pose discrimination task ensures: a) a detailed atomic-level balance between desolvation and other non-bonded interactions, and b) a balance between the protein and generalized energy models. The hydration free energy recapitulation task regularizes solvation parameters in the same type of data as in the protein energetic model^15^.

The *lattice discrimination* task measures how well a given energy function parameterization discriminates near-native lattice conformations against alternate “decoy” conformations for a set of 870 small-molecules. Discriminative power is measured by *Boltzmann probability metric*, which measures the average probability of selecting near-native conformations^15^ with variable definitions of “near-natives” of crystal RMSD of 1, 2, 4, 6 Å. The temperature factor (k_b_T) is defined as 0.1 times the gap between 5 percentile and 95 percentile energy values. Crystal RMSD is measured considering the asymmetric unit and all symmetry mates within 12 Å. Two values are computed and averaged: i) the Boltzmann probability for a set of 100 pre-selected conformations (native and non-native) which are only scored; and ii) the Boltzmann probability for a set of 20 pre-selected conformations that are minimized with the current energy parameters. These decoys are selected at the beginning of dualOptE with initial parameter set and always included at least one sub-Angstrom structure with the lowest energy.

The *atomic geometry matching* task measures the Kullback-Leibler (KL) divergence in the distribution of atomic geometries (non-bonded distances and torsion angles) optimal for an energy parameter set against statistics collected from the *extended training set* of ~4,000 small molecules. Atomic geometry optimal for a parameter set is collected from minimized structures of predicted crystals (see *lattice discrimination task* above) individually for each type of atomic distance and torsion angle.

The *ligand pose discrimination* task measures the Boltzmann probability of selecting near-native ligand pose against a pool of pre-sampled protein-ligand complexes. Pre-sampled complex set comprises both false and near-native poses for 215 various complexes^27^, none of which overlap with any of the targets in our ligand-docking benchmarks. At the beginning of a dualOptE run, 30 conformations with lowest energy (including at least one with ligand RMSD < 1Å with lowest energy) are chosen for each complex. At each cycle of dualOptE, each complex is minimized with the current energy parameterization (fixing the receptor conformation for efficiency), and receptor-ligand interface scores are collected. The Boltzmann probability is measured using the same criteria as in the lattice discrimination task.

The *hydration free energy recapitulation* task measures how well a solvation parameter set recapitulates experimental hydration free energy values of various small molecules, using a dataset of 643 small molecules^28^. The hydration free energy of a molecule is calculated by summation of polar (dG_polar_) and non-polar (dG_nonpolar_) contributions to the total solvation free energy, each of which are estimated as:

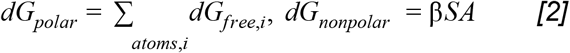

Here, SA is the surface area of the molecule, and *α* and *β* are weighing factors on each term. These weighing factors are determined by least-square-fit of this equation to experimental free energy values of amino-acid analogues^29^ by taking dG_free,i_ values determined for protein atom types. Net agreement is measured as the sum of absolute errors in calculated values (in kcal/mol) over these 643 molecules.

Finally, we validated the parameters on a list of thermodynamic liquid properties (density and heat-of-vaporization) shown in **Fig S5**.

### GALigandDock: A genetic algorithm based Ligand docking method in Rosetta

We developed a new small-molecule docking tool within Rosetta, *GALigandDock*, that enables fully automated on-the-fly sampling of both receptor and ligand conformational space. This docking tool iteratively evolves a pool of protein/ligand complex conformations against *RosettaGenFF*. It makes use of several key features broadly utilized in the ligand docking field: a motif-guided search for initial ligand placements, genetic algorithm optimization, and a grid-based energy precomputation. Each run takes 5 to 20 CPU-minutes depending on system size and receptor flexibility setup.

#### Overview of Docking method

*GALigandDock* accepts a single complex structure as input, and searches for a pool of structures optimal for our generalized energy model through a genetic algorithm. While its basic algorithm adopts broadly accepted ideas in the ligand docking field, several unique features are also utilized. Only DOFs describing the ligand conformation (including 6 rigid body DOFs and DOFs describing rotatable torsions) are encoded into genes. If receptor flexibility is used, additional precomputation of the energy values of flexible parts is carried out; those “implicit” DOFs are optimized on-the-fly in their internal coordinates for every structure generated during genetic algorithm. The protocol starts with optimizing receptor side-chains and their protonation states at apo-state (except for self-docking). Then a subset of the initial pool was generated by *motif-guided ligand conformation search* (see below) portion of which varying between 50~70% depending on the number of possible motif match combinations (more the higher portion), and the rest from randomized genes.

At every iteration in the genetic algorithm, a gene undergoes either mutation (20% chance) or crossover (80% chance) with a randomly selected gene. For every generated conformation, receptor side-chains are optimized by a Monte Carlo (MC) search in discrete rotameric space followed by quasi-Newton minimization in all torsions including those in the ligand, repeating this twice by first ramping LJ repulsion scale from 0.1 at the first cycle to 1.0 at the second cycle. Both MC and minimization are efficiently carried out using a *3D grid representation of energy* (see below). In the 10,000 steps of MC search, one-body and two-body energies of rotamers precalculated at the beginning are utilized^30^. Input rotamers possess constant bonuses of −2.0 kcal/mol in their one-body energies in order to prevent drifting away too much from the input. The 100 “parents” and 100 “children” are then pooled and trimmed to the lowest-energy 100 not closer than 1Å to one another; these 100 serve as the next generation’s “parents.” After 10 iterations, the top 20 structures are further side-chain optimized and backbone- and sidechain-minimized using the ungridded (continuous) energy. A single structure having the lowest complex energy is taken as a single representative.

*GALigandDock* supports a fully automated receptor flexibility logic. Initially, an ellipsoid is constructed around the input ligand conformation. Moments of inertia are computed and are scaled by the half size of the ligand box; if the moment of inertia along an axis is < 0.1 it is increased to 0.1 (for planar molecules). All protein residues whose average side-chain position overlaps this ellipsoid are assigned as flexible. On average 9.8 side-chains are assigned as flexible in the cross-docking benchmark set. There could be a possible caveat that assignments can be sensitive to initial ligand placement for an elongated ligand.

The simulation is repeated 5 and 16 times with median runtime running single simulations of self-docking and cross-docking are 8.5 and 19.7 minutes, respectively. Multiplying by the number of repeats made per task, median core-hours per target in this study are 0.7, 5.3 hours, respectively. All the computational performances were benchmarked in Intel E5-2650 v2 2.2 GHz processors. Examples of running *GALigandDock* can be found in **Supplemental Data**.

#### 3D grid representation of energy

*RosettaGenFF* is represented in 3D energy grids around the ligand pocket, which allows over 10-fold speed-up of docking simulations^31^. For each atom type in the ligand, a per-atom “energy field” is computed on a 0.25 Å grid in a cubic box covering the pocket. The size of the cubic box is allocated depending on the maximum heavy atom distance from center-of-mass of the ligand (*r_max_*), more specifically, as

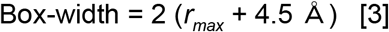

This results on average 24 Å of cell dimensions in a cubic box. The energy field summarizes the interaction of all *rigid* receptor atoms to an atom at a particular grid point, allowing ligands to be scored against the grid without explicit enumeration over individual atomic pairs. 3D spline interpolation is used to compute and minimize off-grid points. Flexible side-chains do not contribute to grid energetics.

Our full *RosettaGenFF* energy model was reproduced in a grid representation. Special treatment was required for several orientation dependent terms (as graphical illustrations shown in **Fig 2a**) highlighted below. For each of attractive and repulsive contributions to E_Lennard-Jones’_, and the isotropic portion of E_Implicit-Solvation_(see Eq. 1), separate grid tables were generated for each of the flexible atom types present. Grid table for the Coulombic term is unified into one representing the electric field. For the orientation-dependent hydrogen-bond term, the sparsity of interactions was exploited: a 3D hash table of receptor donors and acceptors was precomputed, allowing hydrogen bonds to be quickly identified and scored exactly with full consideration of orientation. For the orientation-dependent solvation terms, we could not exploit similar sparsity. Instead, these were approximated as the sum of two isotropic terms per-atom: one based on the atom position, and one based on a “water-binding” virtual position.

**Fig 2.**
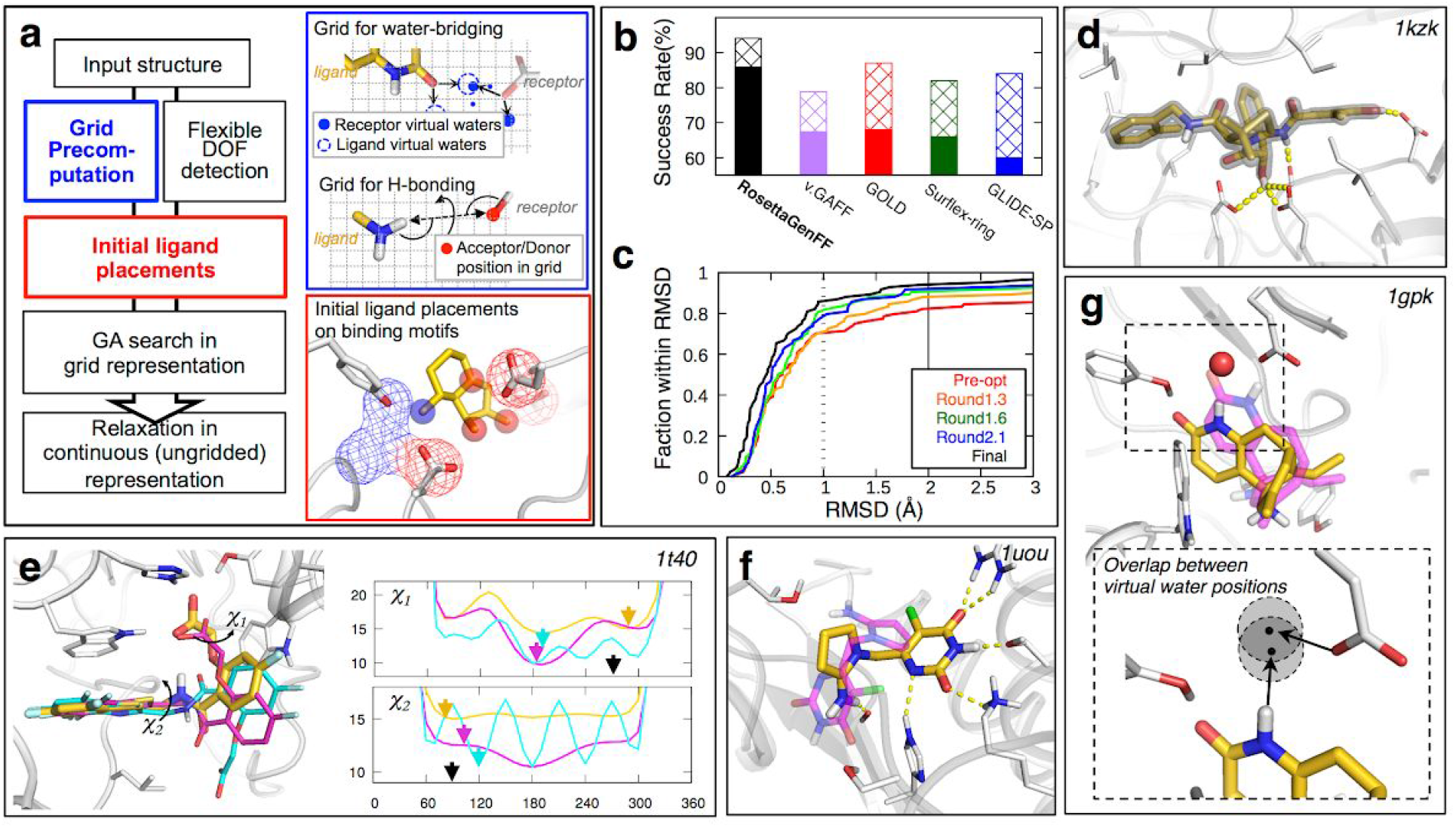
Improved force field leads to more accurate small-molecule pose predictions. **a)** Schematic description of Rosetta *GALigandDock* protocol. Graphical illustrations of steps highlighted in colors are shown in insets with corresponding colors (details in **Methods**). **b)** Self-docking results using *RosettaGenFF* and *GALigandDock* compared to the best reported results using state-of-the-art docking tools^44–46^ tested on the Astex diverse set^32^. Success rate as assessed by ligand RMSD < 1 Å and < 2 Å in solid and patterned bars respectively. “v.GAFF” stands for *GALigandDock* runs using GAFF instead of *RosettaGenFF*. **c)** Success rate using energy parameters from different stages of optimization; *preopt*, pre-optimized version; *round1.3*, after 3rd iteration; *round1.6*, after 6th iteration; *round2.1*, after first iteration of solvation parameter optimization (7th iteration in total); *RosettaGenFF*, the final parameter set. **d-g)** Examples of structures with highly accurate docked ligand poses. Ligand models are colored in gold for *RosettaGenFF*, in magenta for GAFF, and cyan for RosettaGenFF with the torsion term replaced with ChemPLP used in GOLD^47^, respectively. **d)** A high accuracy prediction with ligand RMSD of 0.2 Å for a molecule with 12 rotatable internal torsions. **e)** An example showing the importance of balance between torsion angle preference and non-bonded interactions, 1t40. Right panels, ligand internal energy profiles as a function of *χ*_1_ and *χ*_2_ torsions are shown for different energy functions. The torsion angles in the predicted pose are indicated by arrows using the color scheme of d-g), the values in the crystal structure are indicated by black arrows. **f)** An example highlighting the importance of orientation-dependent hydrogen bonding term, 1uou. *RosettaGenFF* prefers a ligand pose with rich hydrogen bonding (RMSD 0.3 Å) while GAFF prefers one with more solvent exposure (RMSD 5.4 Å). **g)** An example of benefit provided by orientation-dependent water-bridging energy term^20^. Crystal water depicted in red sphere is not modeled explicitly in docking simulation, but still the water-bridging term gives a bonus when virtual water sites overlap (bottom inset) leading to RMSD 0.2 Å prediction; best pose by GAFF lacking this term clashes with this water position (RMSD 1.4 Å).

Comparing exact to grid-computed energies, we see a Pearson correlation of 0.95, with most of the error coming from the orientation-dependent solvation terms (0.84 Pearson correlation).

#### Motif-guided ligand conformation search

It is critical for the genetic algorithm to start with a pool of genes that are promising but are also diverse. In initial testing, we found that fully randomized starting conformations had difficulty with ligands making hydrogen bonds deep inside the pocket. Therefore, a motif-guided placement strategy was applied for about 2/3 of our starting pool (50-70 models out of 100, with greater numbers for receptors with many pocket hydrogen bond donors or acceptors). All non-solvent-exposed hydrogen bonding sites in the receptor are identified, and “ideal waters” are built from these sites representing possible hydrogen-bonding ligand atom positions. These waters are clustered using a 4Å radius, and the *N* clustered motifs with best sum-of-grid-scores having at least two members are selected. N is always between 5 and 15, if too large, stricter criteria for solvent exposure are used; if too small, 1-member “clusters’’ are also considered. Groups of hydrogen-bonding atoms in ligands are defined as ligand motifs with the same clustering criteria. Motif matching and optimization of ligand conformation is then carried out for every possible pair combination of *M* receptor-to-ligand motif matches (M <= 70): for each motif-match, we first translate the ligand to the position where the center of mass of selected motifs overlap, followed by random sampling of ligand orientation and torsion angles; the best after 200 random trials is then minimized against the grid with distance restraints favoring designated motif match. Maximum 70 ligands conformations are generated, each from a unique motif match, prioritizing those matches with higher sum-of-grid-score. if M <= 50, search on the matches with higher sum-of-grid-score are repeated until 50 conformations are generated.

#### Ligand Docking Dataset

We used Astex diverse^32^ and non-native^33^ sets for self- and cross-docking benchmarks, respectively. Ligand protonation states are fixed as provided in the original mol2 files. When testing ligand docking using a conformation directly built from its chemical connectivity (i.e. SMILES string), its initial conformation was generated by CORINA^34^ with a few corrections to the protonation states in the output: carboxylic acids and protonated phosphates are deprotonated (as protonation overly preferred by CORINA). We then further optimized the geometry with AM1 calculation^35^ using Antechamber in AMBER suite. An extended self-docking benchmark set consisting of 212 complexes was brought from a subset of previous work^27^ (list in **Supplemental Data**).

## RESULTS

### Small molecule crystal lattice discrimination

We evaluated the parameterized force field by predicting the crystal structures of 516 small molecules from CSD not used in training. We define “success” as selecting a crystal lattice less than 1 Å RMSD to the experimentally observed lattice as one of the 10 lowest energy structures (crystal lattice prediction is a quite non-trivial challenge)^5^. We compared performance to the generalized Amber force field (GAFF)^1^, which, like our energy function, is sufficiently fast that it can be used for drug discovery studies^36,37^. GAFF had an advantage over other such force fields, in addition to its popular and broad usage, for validation as it could be readily implemented in Rosetta for direct comparison to *RosettaGenFF*; note that GAFF was not optimized using small molecule crystal data. On the validation set, *RosettaGenFF* outperformed GAFF in both the Boltzmann weight^15^ of the observed crystal structure in the population of sampled structures, and in the success rate with the definition aforementioned (**Table 1**; 58% by *RosettaGenFF* compared to 30% by GAFF). Two classes of functional groups stand out when the performances of *RosettaGenFF and* GAFF were compared on a per-group basis (**Fig 1c, Fig S1-2**). Improved results were obtained for *polar conjugating groups* (e.g. esters or aryl-nitros) likely because of the improved balance between torsional and non-bonded energy parameters leading to better transferability across different chemical contexts. Improved results with *hydrogen-bonding groups* are likely due to the explicit treatment of the orientation dependence of hydrogen-bonding in *RosettaGenFF*, an improvement over the GAFF isotropic point-charge model.

**Table 1.** Performance on various training tasks following optimization

|  | Tasks | Measure | Pre-optimize | Optimized | GAFF |
| --- | --- | --- | --- | --- | --- |
| Training | Small-molecule Xtal docking | Boltzmann Probability<sup>1)</sup> | 0.470 | 0.652 | - |
|  |  | Success rate (%)<sup>2)</sup> | 39.3 | 63.6 | - |
|  | Dihedral distribution | Mean KL-divergence | 0.355 | 0.225 | - |
|  | Distance distribution |  | 0.173 | 0.162 | - |
|  | Ligand pose prediction | Boltzmann Probability<sup>1)</sup> | 0.529 | 0.610 | - |
|  | Hydration free energy | Error (kcal/mol) | 6.4 | 2.0 | - |
| Validation | Small-molecule Xtal docking<sup>3)</sup> | Boltzmann Probability<sup>1)</sup> | 0.321 | 0.640 | 0.386 |
|  |  | Success rate(%)<sup>2)</sup> | 23.5 | 58.3 | 29.9 |

- 1) Boltzmann probability selecting near-native structure against non-native ones<sup>15</sup>. Values reported are values averaged over 4 criteria of near-native definitions, each corresponding to 1,2,4,6 Å of crystal-interface RMSD measured on the central asymmetric unit and all symmetry mates within 12 Å. - 2) Success defined as any sub-Angstrom structure within 10 lowest energy structures sampled. - 3) Compared against a common set of 430 molecules having at least 5 of sub-Angstrom structures sampled in all cases.

Even with explicit fitting to lattice data, there is clear room for improvement in our energy model. Proper consideration of polarization effects, in particular a general and higher-level description of anisotropic hydrogen bonding and orbital conjugations in torsions, is an important future direction. Methods with proper treatment of polarization effects -- such as density functional theory (DFT) methods or polarizable force fields^5^ -- achieve better performances in crystal structure prediction, with success rates of 70-80%. However, such methods are too slow for large-scale drug discovery problems. A force field with similar efficiency to ours by Broo et al^38^, specifically designed for crystal lattice docking, performed similarly to ours (50% success rate on their own test set, compared to 51 % with *RosettaGenFF* on the same set). One possible future avenue for improvement would be introducing off-atom charges^39,40^.

### Small molecule docking with *RosettaGenFF*

We investigated the use of *RosettaGenFF* for small-molecule docking calculations using the newly developed Rosetta *GALigandDock*. Accurate ligand pose prediction through molecular docking is of great importance in drug discovery as it provides detailed information about interacting protein residues, and is critical for accurate estimation of relative or absolute binding free energy of potential binders^6,41,42^. A unique strength of our approach comes from the grid-representation of water-bridging effects^20^ and hydrogen bonding in *RosettaGenFF*, both are orientation-dependent and are identified as important features in ligand/protein energetics.

We first tested the new energy function and docking method on 85 complexes from the Astex diverse self-docking set^32^ keeping the protein backbone and side-chain fixed. *RosettaGenFF* incorporated into *GALigandDock* produced lowest energy models with a median RMSD of 0.45 Å, with success rates of 86/94% predicting ligand conformations within 1/2 Å RMSD of the crystal conformation, and 31/56% within 0.3/0.5 Å of the crystal structure, respectively. This high success rate and atomic accuracy suggests that the new energy model successfully identifies both the correct minima in large conformational space as well as precise geometry within the energy basin (**Fig 2d-g**). When docking calculations were performed on a set of ligand conformations directly built from scratch using chemical connectivity (i.e. SMILES)^43^, results are slightly worse, giving a median RMSD of 0.59 Å and success rates of 80/92% using 1/2 Å criteria. Despite failures arising from input ligand structures not well handled in our docking simulations (**Fig S3**), in both cases the results were better than the other methods on the same set (**Fig 2b**)^44–46^. The combination of *RosettaGenFF* and *GALigandDock* on an extended docking set of 212 complexes -- non-overlapping with any target in other protein-ligand training/test sets -- again showed a performance superior to GOLD^47^ with 10% difference in success rates (**Fig S4**).

We then repeated the test with variants of the energy function. A clear improvement was observed (**Fig 2c**) throughout the course of optimization of *RosettaGenFF*. This alone is a somewhat surprising result as the docking benchmark is quite different from the crystal structure training set (contribution from ligand-pose discrimination test used in training was very minor). We also tested the same docking benchmark i) taking GAFF energy parameters or ii) replacing the torsion term into the empirical torsion term used in GOLD, while keeping the energy model on the receptor unchanged in both tests. Poorer performance was obtained for both tests (**Fig 2e-f**).

We next tested the effectiveness of our energy model and docking protocol on the more realistic *cross-docking* problem, in which compounds are docked onto independently determined structures. *GALigandDock* allows any residue that can potentially interact with the ligand to sample alternative backbone and side-chain conformations, resulting in as many as 20 pocket residues to be optimized along with the ligand conformation. This flexibility is enabled by the ability of the underlying Rosetta protein force field^15,21^ to model the energetics of protein conformational changes, and Rosetta’s tools for side-chain conformational sampling and energy minimization^30^. We tested cross-docking performance on the *Astex non-native* set^33^, a standard benchmark set consisting of 1,112 protein-ligand complexes. On this set, *RosettaGenFF* incorporated into *GALigandDock* achieved a median RMSD of 0.86 Å with success rates of 52/74% (using the criteria of ligand RMSD within 1/2 Å, respectively). This is an over 10% improvement in success rate over any previously reported study reported to date on the set^27,33,48–51^ (**Fig 3a**).

**Fig 3.**
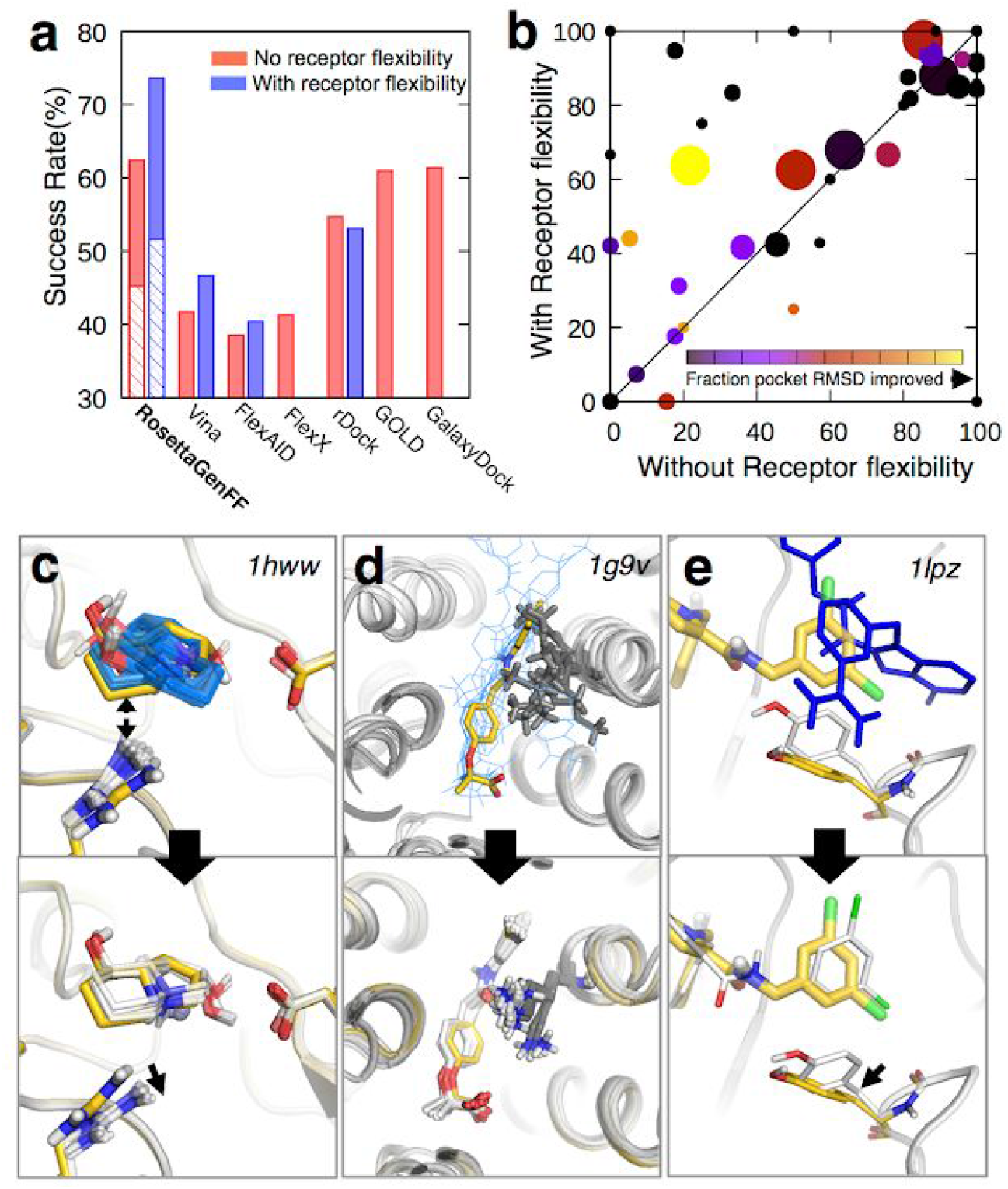
Incorporating receptor flexibility improves cross-docking. **a)** Success rates in cross-docking benchmark for various methods^27,33,48–51^ tested on Astex non-native set^33^. Blue and red bars represent results from docking runs with and without receptor flexibility, respectively; solid and patterned bars show results by two criteria, ligand RMSD < 2 Å and < 1 Å, respectively. Sub-Angstrom success rates are not achieved with other methods. **b)** Per-protein cross-docking results compared between with (Y-axis) and without receptor flexibility (X-axis). Size of points represent number of alternative protein conformations from largest (>50) to smallest (<10); colors represent fraction of conformations with pocket RMSD improved or unchanged by flexibility, from 0.0 (black) to 0.8 (yellow). **c-e)** Examples in which flexible docking improves prediction. Top and bottoms panels are predictions without and with receptor flexibility, respectively. Crystal poses shown in gold, predicted ligand poses starting from multiple receptor conformations in blue (top panels) or white (bottom panels). **c)** 1hww, clash with arginine is relieved, increasing fraction of predictions within sub-Angstrom accuracy from 18% to 95%. **d)** 1g9v, rotameric search on lysine helps increase sub-Angstrom accuracy from 22% to 60%. **e)** 1lpz, starting conformation from PDB ID 1f0s, backbone flexibility allows to correct the orientation of tyrosine leading to ligand RMSD 0.9 Å (10.4 Å without receptor flexibility)

Comparing these results to those without receptor flexibility showed that improvements in ligand pose accuracy primarily came from complexes in which pocket side-chain accuracy also increased (**Fig 3b**); relieving small clashes (**Fig 3c**), correcting wrong sidechain rotameric states (**Fig 3d**), and modeling small backbone conformational changes (**Fig 3e**); note that all of these were achieved by fully automated flexibility annotations. Of 277 complexes with initial models having relatively accurate backbones (RMSD < 1 Å) but for which rigid-receptor docking failed, about half (139) were successfully docked (ligand RMSD < 2 Å) following incorporation of receptor flexibility. The balance between protein and non-protein energetics is clearly important for flexible backbone docking^40^.

## DISCUSSIONS

The small molecule docking results described in this paper demonstrate the power of using prediction of small molecule crystal lattices, a new source of data, to drive energy model parameterization for accurate molecular docking studies. *RosettaGenFF* outperforms previously reported approaches when tested on a range of structure-based drug discovery applications. In the context of the functional forms used, the current energy model may be quite close to optimal for protein/ligand docking: when any of energy components or flexibility was varied from current implementation, around 10% worsening was observed in cross-docking (**Table S2**). Avenues for future improvement include improving the underlying physical model, for example: a) introducing an efficient polarizable and/or multipole electrostatic model^52,53^ and b) additional bonded terms for ring systems^54^. A large amount of small molecule crystal data that was not used in this study could be utilized for this further development, which could improve coverage in chemical diversity. Incorporation of quantum chemistry data during training could further improve the model, particularly for binding free energy calculations.

The combination of *RosettaGenFF* and *GALigandDock* can be readily applied to high-precision virtual screening problems; to this aim, it will be beneficial to enhance its computational efficiency for high-throughput predictions. Both computational advances, such as GPU-accelerated calculation, and algorithmic improvements, such as a “competition-style” model where ligand identity can change along with ligand conformation in the genetic algorithm, should improve run-time, allowing for screening against very large ligand libraries.

