## Supplemental Info for "Learning a force field from small-molecule crystal lattice predictions enables consistent sub-Angstrom protein-ligand docking"

Table of Contents

1. Supplemental Tables
    - Table S1. List of atom types in RosettaGenFF
    - Table S2. Performance of docking benchmarks introducing variations in energy or protocol

2. Supplemental Figures
    - Fig S1. Per-group decomposition of Fig 1c
    - Fig S2. Examples of small-molecule crystal structure prediction highlighting difference between energy models
    - Fig S3. Examples of docking failures starting from SMILES string
    - Fig S4. Benchmark result on extended self-docking set
    - Fig S5. Prediction of thermodynamic properties with RosettaGenFF
    - Fig S6. Reproducibility of GAFF by in-house version

3. Supplemental Data
    - List of PDB IDs in the extended docking set
    - Example for generating a Rosetta parameter file containing generic atom types
    - Example input for small molecule crystal structure prediction
    - Example input for running GALigandDock

**Supplemental Tables**

**Table S1.** List of atom types in RosettaGenFF.

| Element | Name | Hybridization state | Subtypes[1)] | H-bond behavior[2)] | Representative functional group | Other functional groups |
|---|---|---|---|---|---|---|
| C | CS | sp3 | 0,1,2,3 | | | |
| | CD | sp2 | 0,1,2 | | | |
| | CR | aromatic | | | | |
| | CT | sp1 | | | | |
| | CSp | sp3, bound to N/O/S | | | | |
| | CDp | sp2, bound to N/O/S | | | | |
| | CRp | Aromatic, bound to N/O/S | | | | |
| | CTp | sp1, bound to N/O/S | | | | |
| N | Nam | sp3 | | D | Ammonium | |
| | Nam2 | sp3 | | A,D | Amine (with a lone pair) | |
| | Nad | sp2 | | D | Primary/2ndary amide | |
| | Nad3 | sp2 | | D | 3rd amide | |
| | Nin | aromatic | | D | Indole | Pyrrole, |
| | Nim | aromatic | | A | Imidazole | Pyridine, Pyrimidine, Pyrazole |
| | Ngu | sp2 | 1,2 | D | Guanidinium | Aniline |
| | NG1 | sp1 | | | Nitrile | |
| | NG2 | sp2 | 0,1,2 | | Azo | |
| | NG3 | sp3 | | D | - | |
| O | Ohx | sp3 | | A,D | Hydroxyl | Carboxylic acid |
| | Oet | sp3 | | | Ether | Ester, Acetyl |
| | Oal | sp2 | | A | Aldehyde | Carbonyl, Carboxylic acid |
| | Oad | sp2 | | A | Amide | |
| | Oat | sp2 | | A | Carboxylate | |
| | Ofu | aromatic | | | Furan | |
| | Ont | sp2 | | | Nitro | |
| | OG2 | sp2 | | A | - | |
| | OG3 | sp3 | 0,1 | A | Oxime | |
| H | HC | Bound to C | | | - | |
| | HR | Bound to CR | | | - | |
| | HO | Bound to O | | polarH | - | |
| | HN | Bound to N | | polarH | - | |
| | HS | Bound to S | | | - | |
| | HG | Bound to else | | | - | |
| S | Sth | sp3 | | | Thiol | |
| | Ssl | sp3 | | | Sulfide | |

| | | | | | | |
|---|---|---|---|---|---|---|
| | SG2 | sp2 | | | Sulfone | |
| | SG3 | sp3 | | | Sulfate | |
| | SG5 | sp3d2 | | | | |
| | SR | aromatic | | | Thiophene | Thiazole |
| P | PG3 | sp3 | | | Phosphate | |
| | PG5 | sp3d2 | | | | |
| Halogens | F | aliphatic | | | | |
| | Cl | aliphatic | | | | |
| | Br | aliphatic | | | | |
| | I | aliphatic | | | | |
| | FR | aryl | | | | |
| | ClR | aryl | | | | |
| | BrR | aryl | | | | |
| | IR | aryl | | | | |

1) Defines sub-atom types further specifying the number of hydrogens attached to the heavy atom. Share the same LJ and torsional parameters but differ in solvation parameters.
2) Determines the behavior of Rosetta hydrogen-bonding energy term. A: acceptor; D: donor.

**Table S2. Performance of docking benchmarks introducing variations in energy or protocol**

| **Tests** | **Metrics** | **Variations in Energy** | | | **W/o receptor flexibility** |
|---|---|---|---|---|---|
| | | **RosettaGenFF** | **Pre-optimize** | **w/o orientation terms** | |
| Self-docking | % < 1Å | 85.9 | 70.6 | 80.0 | - |
| | % < 2Å | 94.1 | 82.4 | 89.4 | - |
| Cross-docking | % < 1Å | 51.6 | 45.0 | 43.3 | 45.2 |
| | % < 2Å | 73.6 | 62.6 | 62.6 | 62.1 |
| Virtual screening | AUC | 0.80 | 0.71 | - | 0.77 |

**Supplemental Figures**

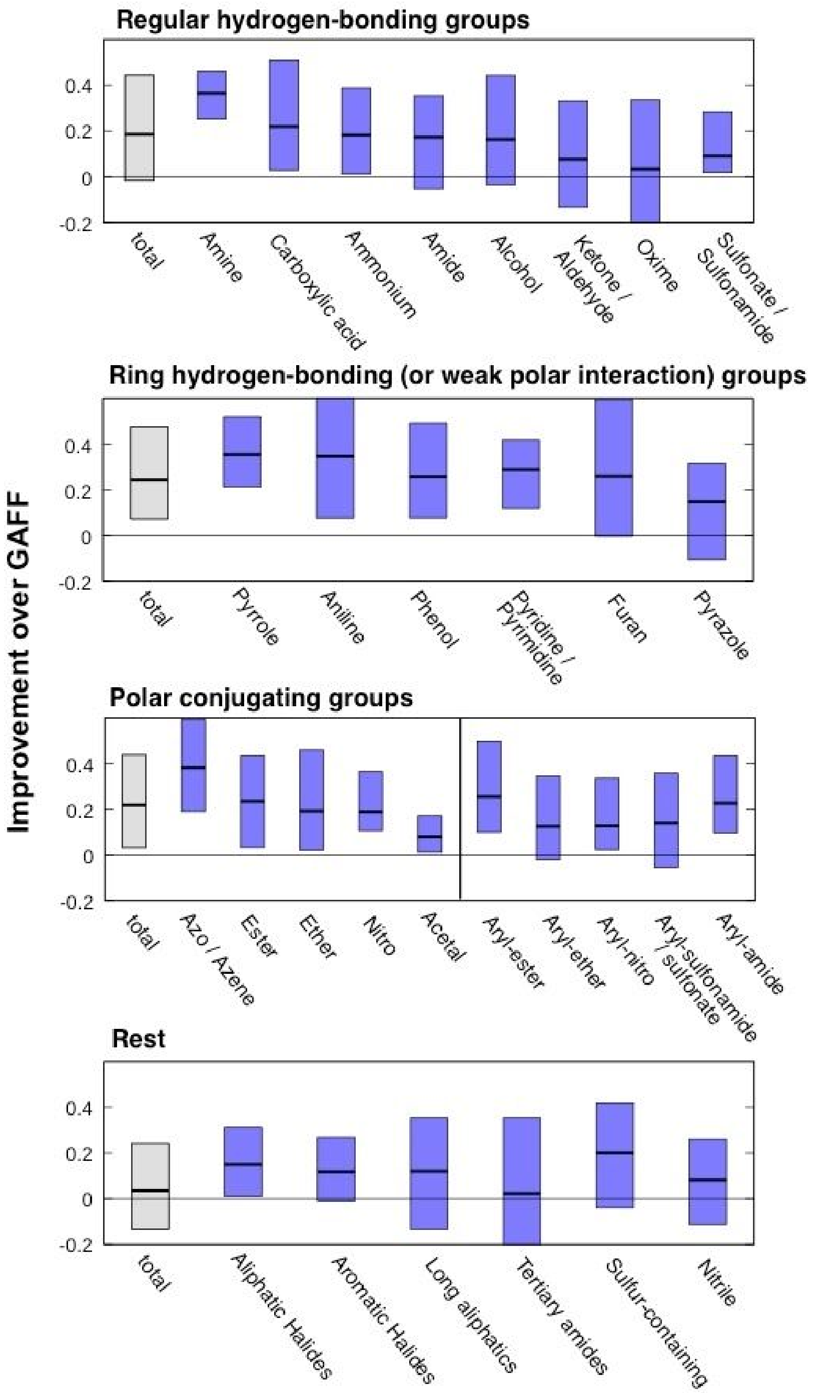


**Fig S1.** Per-group decomposition of Fig 1c**.** Detailed per-functional group analyses on the performance of small molecule crystal structure prediction with RosettaGenFF and GAFF.

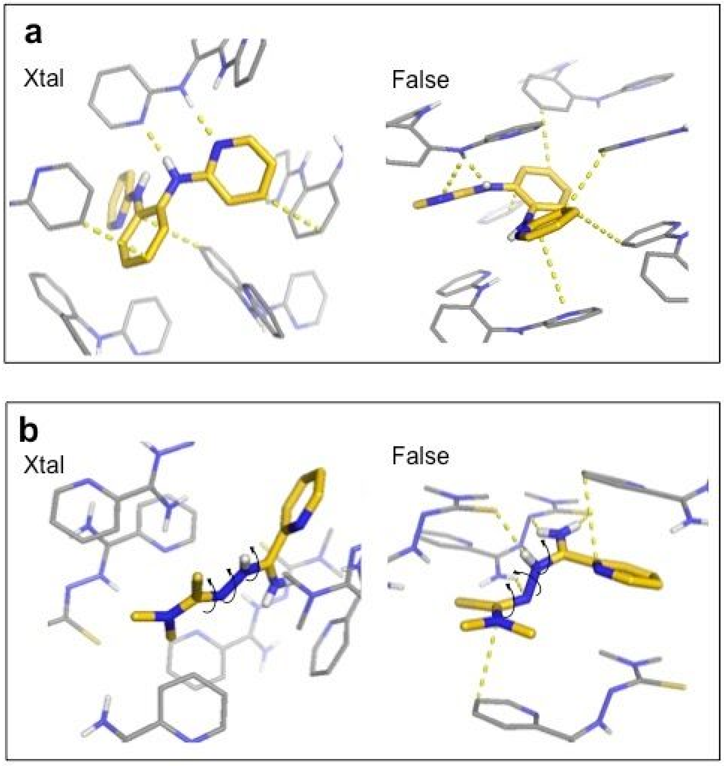


**Fig S2.** Examples of small-molecule crystal structure predictions highlighting differences between energy models: RosettaGenFF, energy minima very close to Xtal conformation, and GAFF, energy minima shown at the right-hand side panels.

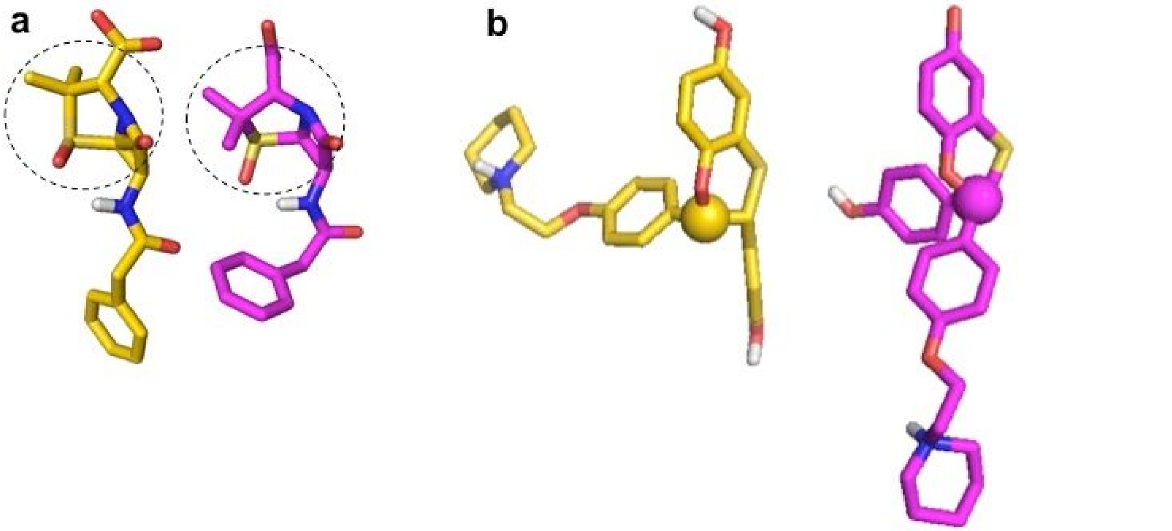


**Fig S3.** Examples of docking failures in Astex diverse set when started from SMILES string: a) 1gm8 and b) 1sj0. Crystal ligand conformation shown in gold, and conformation built from SMILES in magenta.

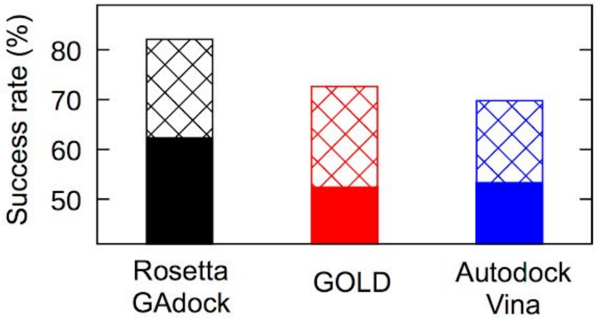


**Fig S4.** Benchmark result on extended self-docking set. Three docking tools are benchmarked against 212 targets independent of protein complexes used for training and other tests in this work. Success rate as assessed by ligand RMSD < 1 Å and < 2 Å in solid and patterned bars respectively. GOLD [1] and Autodock Vina [2] results were collected from local runs. GOLD results were collected by selecting top scoring pose from 25 independent GOLD version 5.7.0 docking runs with default options (including Chem-PLP term). AutoDock Vina results were collected by selecting top scoring pose from 5 independent AutoDock Vina version 1.1.2 runs with default options.

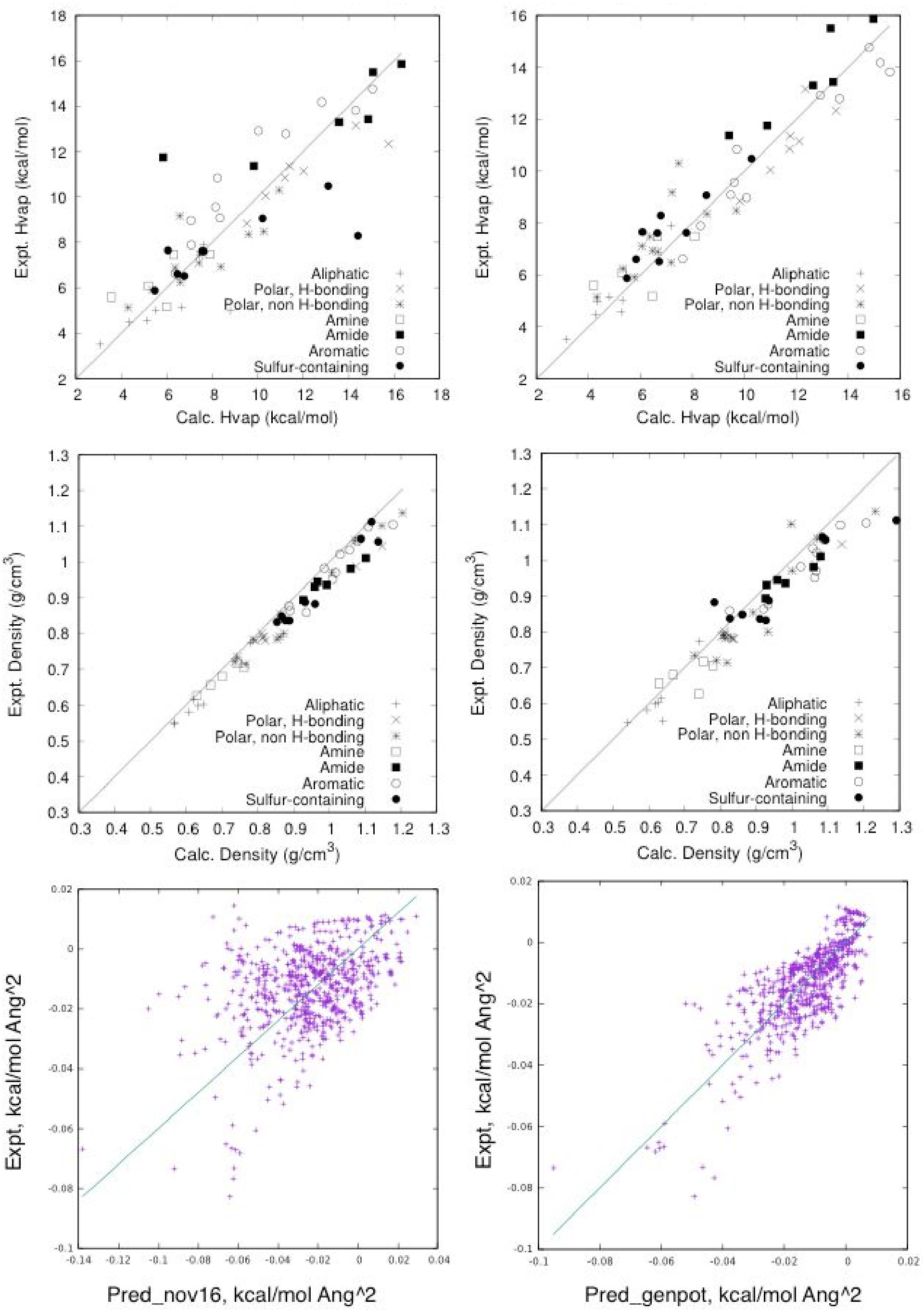


**Fig S5.** Prediction of thermodynamic properties with RosettaGenFF. In all panels, experimental values are shown in y-axis, and calculated values in x-axis. Values predicted by the protein-only energy model and RosettaGenFF are shown in the left and right side, respectively. Top) Prediction of heat-of-vaporization of small molecule liquids brought from Wang et al [3]. Middle) Prediction of density of small molecule liquids brought from Wang et al. Bottom) Prediction of small molecule hydration free energy brought from Mobley et al [4].

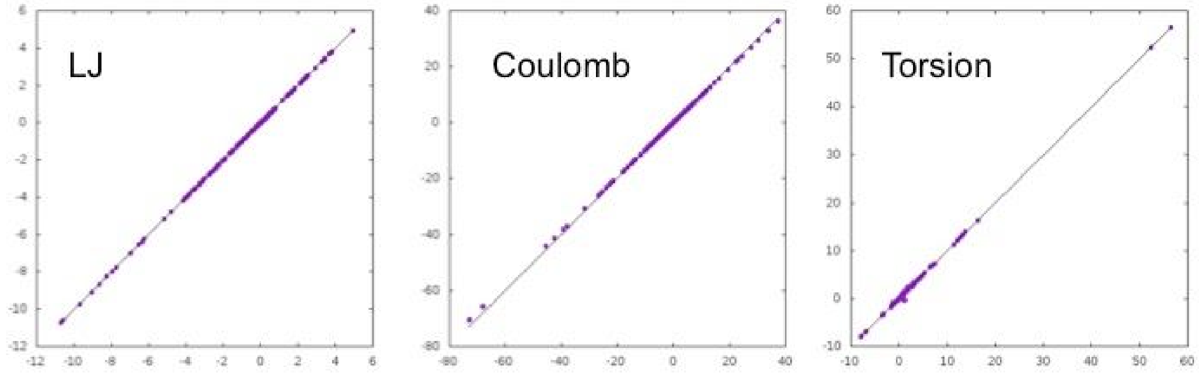


**Fig S6.** In-house version GAFF implementation. GAFF is locally implemented in Rosetta suite to run lattice structure predictions and *GALigandDock*. Point energy calculation results from *sander* in AMBER14 and GAFF version 1.5 are taken as reference for the comparison. Energy values calculated from AMBER14 are compared to those values from our in-house version for a mixture of 330 dimeric and 250 monomeric conformations from 20 different molecules. Lennard-Jones, Coulomb, and torsion terms are reproduced with Pearson correlation of 0.974, 0.985, 0.982, respectively, when tested on 330 dimeric and 250 monomeric conformations of small molecules.

**Supplemental Data**

**1. List of PDB IDs in extended docking set**

1bzy 1duv 1ew9 1f74 1fcz 1flr 1fzq 1gar 1hfs 1hn4 1hvk 1i5r 1ii5 1jyq 1k27 1k6v 1kel 1lst 1lyx 1m7y 1mai 1moq 1njs 1o86 1pme 1pro 1qin 1rmz 1ro6 1t31 1tpw 1ui0 1uj5 1utj 1wc1 1wht 1ws1 1x39 1xjd 1xws 1yei 1zvx 2ccc 2cen 2drc 2epn 2f94 2fgu 2gyi 2hzl 2hzy 2i0a 2i2c 2iko 2jew 2jg0 2oxn 2ptz 2q8h 2ri9 2rkd 2sim 2v3u 2ves 2vj8 2vmd 2vw2 2w8j 2x8z 2xeg 2xii 2xn5 2yfx 2zxg 3a2o 3a5y 3a6t 3ao2 3b1m 3b92 3brn 3bu1 3c2u 3c88 3czv 3d0e 3ebh 3egt 3f7i 3fee 3gba 3hkw 3ies 3ip8 3ipq 3kek 3kjd 3kv2 3lxe 3lxk 3mi2 3n9r 3nxq 3nzk 3p5o 3pcg 3ppm 3pyy 3re4 3rlb 3rux 3rwp 3s0d 3t1m 3t2w 3tay 3th9 3tzm 3u92 3udd 3uug 3v51 3zm9 4ahu 4ayq 4b0b 4b33 4b6o 4b7r 4b8y 4bps 4bt4 4c1u 4c2v 4clj 4cpy 4d1j 4do4 4dsu 4erf 4f9u 4fai 4gbd 4ghi 4gq4 4gu6 4hf4 4hws 4i71 4iuo 4jx9 4lhm 4ll3 4mpn 4nj9 4p5z 4pft 4pp3 4qge 4ql1 4rpo 4tmn 4ty6 4ual 4up5 4uye 4v24 4w9p 4wov 4xt2 4xy8 4ymx 4z2b 4zji 5am6 5b25 5cp9 5d21 5edl 5eei 5egm 5ehr 5elw 5fh7 5g46 5h1t 5hbs 5hu9 5i1q 5ia0 5l7h 5lom 5m4q 5m77 5my8 5sym 5ta2 5tuz 5vcv 5wqc 966c 1i37 1lnm 1rjk 1w96 1zhy 2vw5 3su3 4nnr 4pmm 4qsu 6b96

**2. Example for generating a Rosetta parameter file containing generic atom types**

Rosetta parameters file for an arbitrary molecule (not supported by default parameter library) with proper generic atom types assigned can be generated through a built-in python script:

```
 > $ROSETTA/source/scripts/python/public/generic_potential/mol2genparams.py \
 -s [your.mol2] (or -l [text file containing a list of mol2 files])
```

Please note that the current script supports “mol2” format only.

**3. Example input for small molecule crystal structure prediction**

The Rosetta command line for running single run of MCM crystal lattice prediction through RosettaScripts application is as below:

```
> $ROSETTABIN/rosetta_scripts -parser:protocol crystdock.xml \
    -parser:script_vars sg=$sgin random=$random \
    -s $pdb \
    -nstruct $nstruct \
    -gen_potential -in:file:fullatom -crystal_refine -overwrite \
    -fa_max_dis 10.0 -elec_max_dis 10.0 \
    -extra_res_fa LG.params \
    -out:file:silent output.out -out:file:silent_struct_type binary
```

with an input XML file “crystdock.xml” as below:

```
<ROSETTASCRIPTS>
  <SCOREFXNS>
    <ScoreFunction name="fullatom" weights="genpot_condensed" symmetric="1">
```

```
    <Reweight scoretype="coordinate_constraint" weight="1.0"/>
   </ScoreFunction>
  </SCOREFXNS>
  <MOVERS>
   <DockLatticeMover name="crystdock" scorefxn="fullatom"
        trans_step="0.5" rot_step="2.5" chi_step="2.0" ncycles="50"
        purturb_init_unit_cell = "0" slide_lattice = "1" spacegroup="%%sg%%"
        randomize="%%random%%" rand_init_pos = "%%random%%"
        rand_init_chis = "%%random%%"
        rand_init_cell = "1" min_init_chis="1" perturb_chi="1"
        perturb_lattice_scale = "1"
        verbose="1" min="1" final_min="1" pert_ucelldim_step = "0.2"
        pert_ucellangle_step = "5" contact_dist = "16" init_min_ncycles="5"
        init_score_cut="0" kT="2" recover_low="1"
        occ_min_cutoff="0.3" occ_max_cutoff="2.0" occ_min_cutoff_init="0.4"
        occ_max_cutoff_init="2.0"/>
        constraint_jump="1"/>
  </MOVERS>
  <PROTOCOLS>
   <Add mover="crystdock"/>
  </PROTOCOLS>
  <OUTPUT scorefxn="fullatom"/>
</ROSETTASCRIPTS>
```

$nstruct is the number of structures to be generated; $pdb is a pdb file storing initial conformation of the molecule; “LG.params" is a Rosetta parameter file of the molecule which can be generated by Rosetta parameter generation script (please see above). For each de novo prediction and native crystal refinement, ($sgin, $random) need to be assigned as (“random”,1) and (“input”,0), respectively. A scorefile “genpot_condensed.wts” should be passed to Rosetta, which is,

```
fa_atr 1
fa_rep 0.55
fa_intra_atr_xover4 1
fa_intra_rep_xover4 0.55
fa_grpelec 3.0
fa_intra_elec 3.0
hbond 3.0
NO_HB_ENV_DEP
```

**4. Example input for running GALigandDock**

The Rosetta command line for running single run of GALigandDock through RosettaScripts application is as below:

```
> $ROSETTABIN/rosetta_scripts -parser:protocol liganddock.xml \
    -parser:script_vars runmode=$runmode \
    -holo.pdb -extra_res_fa LG.params \
    -gen_potential \
    -overwrite \
    -mute core basic protocols.relax \
    -crystal_refine -scale_rb 10.0 \
    -score::hb_don_strength hbdon_GENERIC_SC:1.45 \
    -score::hb_acc_strength hbacc_GENERIC_SP2SC:1.19 \
    -score::hb_acc_strength hbacc_GENERIC_SP3SC:1.19 \
    -score::hb_acc_strength hbacc_GENERIC_RINGSC:1.19 \
    -no_autogen_cart_improper
```

with an input XML file “liganddock.xml” as below:

```
<ROSETTASCRIPTS>
 <SCOREFXNS>
 <ScoreFunction name="dockscore" weights="beta_genpot">
  <Reweight scoretype="fa_rep" weight="0.2"/>
  <Reweight scoretype="coordinate_constraint" weight="0.1"/>
 </ScoreFunction>
 <ScoreFunction name="relaxscore" weights="beta_genpot_cart"/>
 <MOVERS>
  <GALigandDock name="dock" runmode="%%runmode%%"
       scorefxn="dockscore" scorefxn_relax="relaxscore" />
 </MOVERS>
 <PROTOCOLS>
  <Add mover="dock"/>
 </PROTOCOLS>
 <OUTPUT scorefxn="relaxscore"/>
</ROSETTASCRIPTS>
```

Three “$runmode”s are currently supported in GALigandDock: “dockrigid” for a rigid-receptor docking; “dockflex” for a flexible-receptor docking (allowing receptor side-chain remodeling and backbone minimization). Ligand conformation is treated as flexible in all these runmodes.

Input structure file “holo.pdb” file should contain the contents of receptor structure first, cofactor/ions (if any), and the ligand molecule at the end. Rosetta parameter file for the ligand molecule can be generated as above; if there is any (non-ion) cofactor in holo.pdb, its parameter file (say, “cofactor.params”) should be generated and passed to the command line with an additional argument “-extra_res_fa cofactor.params”. More details of GALigandDock is provided at:

https://www.rosettacommons.org/docs/latest/scripting_documentation/RosettaScripts/Movers/GALigandDock

and how to prepare inputs for GALigandDock at:

https://www.rosettacommons.org/docs/latest/GALigandDock-Preprocessing
